# Larger, but not better, implicit motor adaptation ability inherent in medicated Parkinson’s disease patients: a smart-device-based study

**DOI:** 10.1101/707208

**Authors:** Ken Takiyama, Takeshi Sakurada, Masahiro Shinya, Takaaki Sato, Hirofumi Ogihara, Taiki Komatsu

**Author notes:** Correspondence should be addressed to K.T.

## Abstract

Generating appropriate motor commands is an essential brain function. To achieve proper motor control in diverse situations, predicting future states of the environment and body and modifying the prediction are indispensable. The internal model is a promising hypothesis about brain function for generating and modifying the prediction. Although several findings support the involvement of the cerebellum in the internal model, recent results support the influence of other related brain regions on the internal model. A representative example is the motor adaptation ability in Parkinson’s disease (PD) patients. Although this ability provides some hints about how dopamine deficits affect the internal model, previous findings are inconsistent; some reported a deficit in the motor adaptation ability in PD patients, but others reported that the motor adaptation ability of PD patients is comparable to that of healthy controls. A possible factor causing this inconsistency is the difference in task settings, which yield different cognitive strategies in each study. Here, we demonstrate a larger, but not better, motor adaptation ability in PD patients than healthy controls while reducing the involvement of cognitive strategies and concentrating on implicit motor adaptation abilities. This study utilizes a smart-device-based experiment that enables motor adaptation experiments anytime and anywhere with less cognitive strategy involvement. The PD patients showed a significant response to insensible environmental changes, but the response was not necessarily suitable for adapting to the changes. Our findings support compensatory or paretic cerebellar functions in PD patients from the perspective of motor adaptation.

## 1 Introduction

Motor adaptation is an essential brain function that modifies motor commands to achieve desired movements in novel situations, such as learning to use new tools or correcting a movement error. A promising hypothesis about motor adaptation is the internal model hypothesis, which considers the cerebellum to play a role not only in predicting future states of the environment and the body but also in modifying the prediction [1]. Appropriate motor commands can be generated through the outcome predicted by the internal model. To investigate the ability to update the internal model, a motor adaptation paradigm is used [2,3]. In this paradigm, the environment changes through artificially applied perturbations. The subjects thus need to update their internal models to achieve the desired movements while compensating for the perturbation.

The cerebellum can play crucial roles in the internal model [1], which has been supported via deficit motor adaptation abilities in cerebellar ataxia patients [4,5]. In addition to the cerebellar ataxia patients, Parkinson’s disease (PD) patients also showed impaired sensorimotor adaptation abilities, such as adapting to 90 degree visuomotor rotation [6] or adapting to three-dimensional arm-reaching movements [7]. PD, the second most common degenerative neurological disease, causes a lack of dopamine neurons in the substantia nigra pars compacta. Due to the impaired motor adaptation ability in PD patients, the internal model can be affected by not only the cerebellum but also other brain regions. The motor adaptation ability in PD patients can provide some hints to deepen our knowledge about the internal model.

To investigate the motor adaptation ability inherent in PD patients, previous studies relied on a constant amount of perturbation that was applied abruptly at a specific time (we refer to this type of perturbation as abrupt perturbation hereafter). To adapt to the abrupt perturbation, subjects tend to rely on explicit strategies or their cognitive abilities [8,9]. For example, when the perturbation was 90 degree visuomotor rotation that caused a 90 degree deviation in the movement angle between the actual and perturbed movements, it was possible to achieve the desired movements by aiming at the 90 degree location distant from a target. With an abrupt perturbation, subjects tend to notice the onset of the perturbation or task switching [9–11]. In the motor adaptation to the abrupt perturbation, it is difficult to determine whether the impaired sensorimotor adaptation in PD patients is caused by their cognitive abilities (i.e., the influence of the task switch) or adaptation abilities.

In contrast to the adaptation to an abrupt perturbation, PD patients showed a compatible adaptation ability with the healthy elderly individuals in responding to a gradually applied visuomotor transformation (we refer to this type of perturbation as a gradual perturbation hereafter) [12,13]. In contrast to abrupt perturbations, a striking feature of gradual perturbations is their difficulty in being noticed [9–11,14,15]. A task switch can thus be related to an adaptation to the abrupt perturbation and not to an adaptation to the gradual perturbation. The influence of the task switch can be a candidate for interpreting the compatible adaptation ability of PD patients with healthy elderly individuals in adapting to only the gradual perturbation.

Additional support for the influence of task switching on motor adaptation in PD patients is the lack of savings in PD patients. Young individuals and healthy elderly individuals show faster learning in relearning trials than in the initial learning trials, such as in the A-B-A paradigm, which is referred to as savings [16—18]. In contrast, PD patients do not exhibit savings [19,20]. Because task switching can lead to savings, the adaptation ability inherent in PD patients can be influenced by task switching.

Because task switching is a candidate in affecting the motor adaptation ability in PD patients, it is necessary to exclude the influence of task switching in detail. Although a gradual perturbation can involve less task switching than an abrupt perturbation, previous studies used 60 degree visuomotor rotation [12] or a 7.8 cm transformation in 10 cm arm-reaching movements in total [13]. These perturbations can involve the influence of task switching because the aiming direction can deviate from the target location even if the gradual perturbation involves more than a 45 degree visuomotor rotation (Fig. 3 in [9]). It is thus still unclear whether the adaptation ability is still compatible between PD patients and elderly individuals because the task switch can be involved in a gradual perturbation with a large amplitude.

Here, we investigated the motor adaptation ability of PD patients while decreasing the influence of the task switch as much as possible. To reduce the influence of the task switch, we relied on a gradually applied perturbation whose existence was noticeable by 1 out of the 82 participants in our previous study [11]; the visuomotor rotation changed by one degree in each trial, and the maximum value of the rotation was 15 degrees. Based on a previous study, task switches are less involved in gradually applied 15 degree visuomotor rotation than in gradually applied 45 degree visuomotor rotation [9]. We demonstrate that the PD patients showed a larger, but not better, motor adaptation ability than elderly individuals and young individuals, rather than a compatible or impaired ability.

In addition to reducing the influence of the task switch, we also decreased the burden to participate in the motor adaptation experiments. Almost all the previous experiments on motor adaptation relied on manipulanda or pen tablet settings. These settings require the subjects to travel to the laboratory, which can be a burden, especially for patients, to participate in the experiments. To minimize this burden, we utilize a smart-device-based experimental setting that is available to conduct motor adaptation experiments anytime and anywhere [11]. Smart-device-based experiments have been proposed to conveniently investigate motor adaptation or visuomotor abilities for flexible applications [21,22]. Our smart-device-based setting has been validated under several conditions and by comparing it to a conventional experimental setting with manipulanda [11]. We refer to our experimental setting as the POrtable Motor learning LABoratory (PoMLab). The PoMLab can decrease the burden for the participants, PD patients, elderly individuals, and young individuals by removing the need to go to a specific place at a particular time. In this study, we demonstrate a larger, but not better, motor adaptation inherent in PD patients using our PoMLab setting.

## 2 Methods

### 2.1 Participants

Fifty-four subjects participated in the current study; their ages and sexes are summarized in Table 1. This study was approved by the ethics committees of the Tokyo University of Agriculture and Technology, Jichi Medical University, and Kakeyu Hospital. The PoMLab experiments were conducted while each participant was seated on a chair and an experimenter was present and close to the participant. The participants provided written informed consent to participate in this study.

**Table 1.**
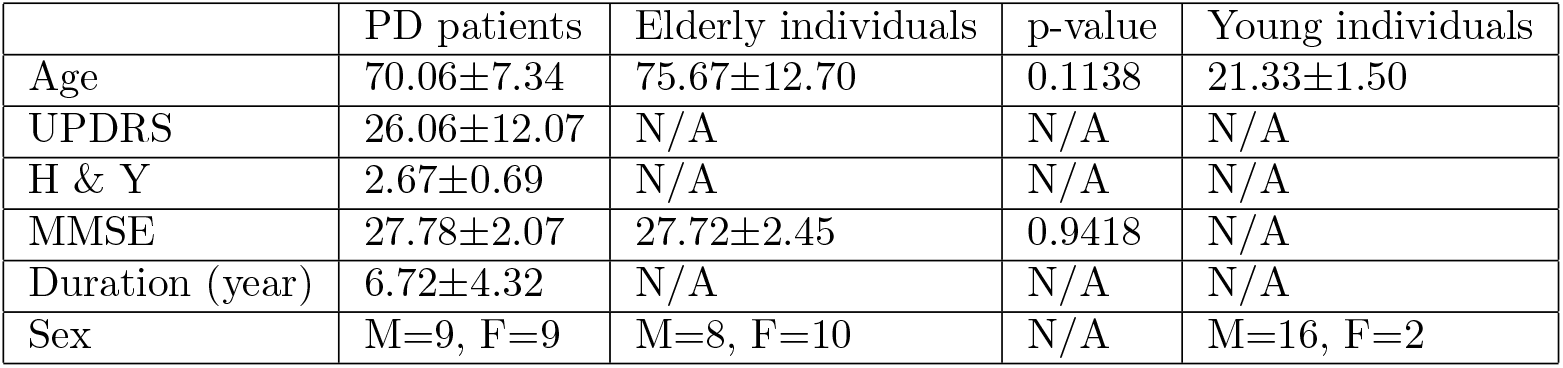
Attributes of the participants. The “p-values” in this table indicate the p-values from a two-sample t-tests between the PD patients and the elderly individuals. “M” and “F” under sex indicate male and female, respectively. All the values in this table denote the means ± standard deviations.

The PD patients were outpatients satisfying the following inclusion criteria. The elderly individuals were inpatients with broken lower-limb bones that did not disturb performance in our study. The young individuals were volunteers. All of the participants were naive to the purpose of the study.

The PD patients were clinically evaluated based on the Unified Parkinson’s Disease Rating Scale (UPDRS) [23] and the adapted version of the Hoehn and Yahr scale (H&Y) [24]. The Mini-Mental State Examination (MMSE) score was used to evaluate cognitive abilities. Table 1 summarizes the ages, UPDRS scores, H&Y scores, MMSE scores, duration, and sex in each group, if available.

### 2.2 Inclusion criteria

PD patients and elderly individuals whose MMSE scores were greater than 22 [no significant difference, two-sample t-test p=0.9418], indicating no severe cognitive decline, were included in the study. Furthermore, the included PD patients were on a medication, resulting in no tremor. There were no musculoskeletal and visual impairments that inhibited performing the required tasks in the current study.

### 2.3 Smart device

We used an Android tablet (Nexus 9, HTC, Taipei City, Taiwan, 2048 × 1536 pixels, 228.25 × 153.68 × 7.95 mm screen size, and 436 g weight) throughout our experiments.

### 2.4 PoMLab application

The PoMLab application was developed using a personal edition of Unity (version 5.2). The PoMLab application is available on our GitHub page (https://github.com/masahiroshinya/PoMLab).

### 2.5 PoMLab settings

The cursor position on the tablet display (*d_x_*, *d_y_*) that was controlled by participants was determined as follows. First, the cursor position in the tablet coordinate system, (*p_x_*, *p_y_*), was determined based on the measured acceleration of the tilting motions in the x- and y-axes of the tablet coordinate system (*a_x_*, *a_y_*) through a low-pass filter; *p_x_* = 0.95*_p_x__* + 0.05 arcsin(*a_x_*) − *o_x_* and *p_y_* = 0.95*_p_y__* + 0.05 arcsin(*a_y_*) − *o_y_*, where *o_x_* and *o_y_* are offsets to determine the initial cursor position in each trial (*o_x_* = 0 and *o_y_* = −30). The *a_x_* and *a_y_* were sampled at 200 Hz. The cursor position in the accelerometer coordinate system was transformed into the position on the tablet display (*d_x_*, *d_y_*) by multiplying by the rotation matrix *R*; (*d_x_*, *d_y_*)*^T^* = *R*(*p_x_*, *p_y_*)*^T^*, where ()*^T^* denotes the transpose of the vector. Without any visuomotor rotation, the cursor position in the accelerometer coordinate system corresponded to that in the display coordinate system. In the *t*th learning trial, visuomotor rotation was applied through the rotation angle p*_t_*. The detailed settings and validations of PoMLab are provided in our previous study [11] and our code is available on GitHub (https://github.com/masahiroshinya/PoMLab).

### 2.6 Experimental procedures

The required task was to tilt the held tablet device appropriately. Corresponding to the tilting motion, the cursor displayed on the tablet moved (a yellow circle with a 4.5 mm radius on the Nexus 9). The participants were instructed to move the cursor toward the visually instructed target (a purple circle with a 4.5 mm radius on the Nexus 9) also on the display in a straightforward manner within two seconds (Fig. 1A). At the beginning of each trial, the subjects needed to tilt the tablet to set the cursor at the initial position in the center of the tablet screen (a blue circle with a 9.0 mm radius on the Nexus 9) for 1 second. After 1 second, the color of the initial position changed from blue to red, and the target appeared. Because the target was displayed for two seconds, the subjects needed to tilt the tablet to hit the target within these two seconds. The cursor, target, and initial position were displayed on the tablet screen, and the cursor moved according to the tilting motion, which enabled the motor adaptation experiments to be conducted solely with the tablet device.

**Fig. 1.**
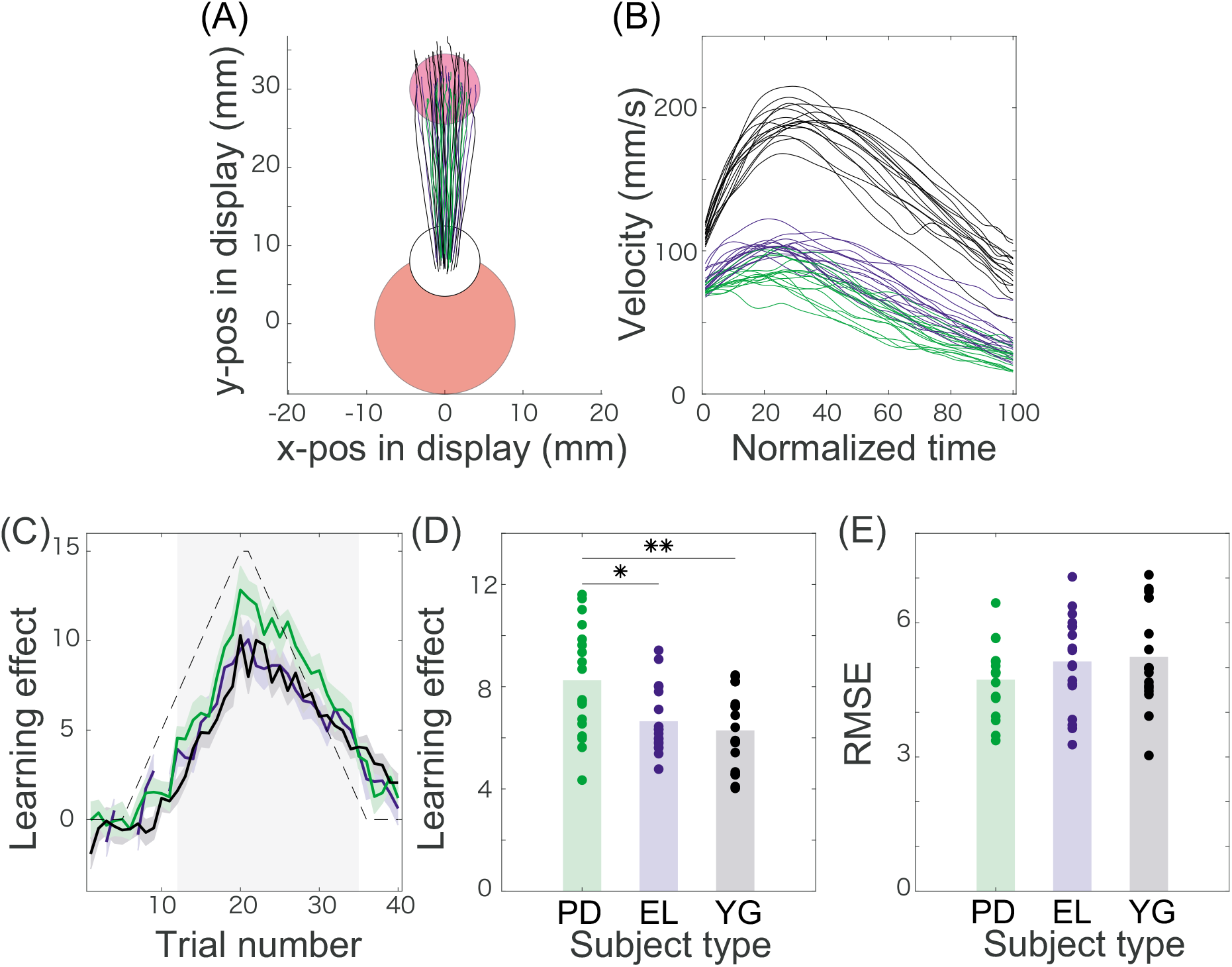
Kinematics and learning curve in the PoMLab experimentso (A): Trajectories displayed on the tablet monitor. The green, blue, and solid black lines denote the averaged trajectories across the PD patients, the age- and MMSE-matched elderly individuals, and young individuals, respectively, in every five trials (N=18 in each group). The red, white, and magenta circles indicate the initial position, the controlled cursor, and the target, respectivelyo (B): Measured velocity. The green, blue, and solid black lines denote the averaged velocities along the y-axis across the PD patients, the age- and MMSE-matched elderly individuals, and the young individuals, respectively, in every five trials (N=18 in each group). (C): Learning curves and the perturbation schedule. The horizontal axis denotes the trial number, and the vertical axis indicates the learning effects or the degree of perturbation (black dotted line). The learning effects were calculated based on the movement angles at the time when the velocity along the y-axis reached its maximal value. The green, blue, and solid black lines indicate the learning effects averaged across the PD patients, the age- and MMSE-matched elderly individuals, and the young individuals, respectively. The green, blue, and black shaded areas indicate the standard error of the mean for the learning effects in each group. The gray shaded area denotes the trial number where learning effects in all the groups are significantly different from zero (t-test p<0.01 [corrected]). (D): Learning effects averaged across the trials denoted in the gray shaded area in panel (C). Each dot indicates the learning effects for each subject. Each bar shows the mean learning effects in each group. * and ** indicate statistically significant differences with p < 0.05 and p < 0.01, respectively (Tukey’s post hoc test following one-way ANOVA). (E): RMSE averaged across the trials denoted in the gray shaded area in panel (C).

The subjects participated in 20 practice trials and 80 learning trials. In the first 20 practice trials, the target position was pseudorandomly set to either 60, 75, 90, 105, or 120 degrees without any visuomotor rotation (90 degrees indicated the 12 o’clock position on the tablet display). In the following 80 learning trials, the target position was fixed at 90 degrees. The learning trials were divided into two parts. In the first 40 learning trials, the subjects experienced gradually increasing and vanishing clockwise (CW) perturbation. In the latter 40 trials, the subjects underwent counterclockwise (CCW) perturbation. Half of the participants experienced the CW perturbation first, and the other half of the participants experienced the CCW perturbation first. No subjects were aware of the existence of the perturbation. The experiment typically took less than 30 minutes.

### 2.7 Evaluation of the learning effects

The learning effects were evaluated depending on the movement angles of the cursor when the velocity on the y-axis reached its peak value (Fig. 1B). The movement trajectories are displayed in Fig. 1A after the movement started. The onsets were detected when the velocity along the y-axis on the display exceeded the mean + 2.5 times the standard deviation calculated in each trial. To avoid evaluating outliers, movement angles in each trial were excluded when these exceeded (15 + *p_t_*) when *p_t_* ≥ 0 or (−15 + *p_t_*) otherwise. There was no significant difference in the number of excluded trials between the PD patient group [3.38±2.96 out of 80 trials] and the elderly group [4.50±4.76 out of 80 trials] (p=0.6053) and between the PD patient group and the young group [1.61±2.20 out of 80 trials] (p=0.2830). There was a significant difference between the elderly and young groups (p=0.0413). We confirmed the invariance of the following results when the exclusion criteria were (*c* + *p_t_*) when *p_t_* > 0 or (*−c* + *p_t_*) otherwise with *c* = 11.5, 12, 13, 14, 16, 20, or 25. When c was less than 11.5, at least one participant in each group showed outliers in the same trial(s) with *p_t_* < 0 and pt > 0. This disturbed the analysis for the age- and MMSE-matched participants in the PD and elderly groups. We thus focused on the case when *c* > 11.5, especially when *c* = 15, throughout this study.

### 2.8 Data analysis

Because outliers were detected in some trials based on the abovementioned criteria, learning effects were averaged across the CW and the CCW conditions in each subject to exclude the effects of the outliers. There were no outliers at the same trials in the CW and the CCW conditions in all the subjects; the averaged learning effects could be reasonably discussed across all the subjects. Because the effects in the CW conditions took positive values and those in the CCW conditions took negative values, we averaged those by multiplying −1 to the learning effects in the CCW condition. If there were outliers at the *k*th trial in a subject in the CW condition, for example, the learning effects at the *k*th trial corresponded to that in the CCW condition.

The averaged learning effects of the ith subject, *x_i_* = (*x*_1,*i*_, …, *x*_40,*i*_), were decomposed into three parameters: the temporal delay Δ_*i*_(Δ_*i*_ ≥ 0), the amplitude *A_i_* (*A_i_* ≥ 0), and the phase *ϕ_i_*. The temporal delay was calculated by temporally sliding the fragments of *x_i_*, *x_i_* (Δ*i*) = (*x*_9+Δ_*i*_,*i*_, …, *x*_32+Δ_*i*_,*i*_), to minimize the squared error from the fragments of the perturbation sequence *p* = (*p*_9_, …, *p*_32_). The squared error between the learning effects and the perturbation is hereafter referred to as the task error. We chose these fragments because the fragments of *x_i_* in the PD group were significantly different from 0 when the trial number was between 12 and 35 (i.e., *x_i_* = (*x*_12,*i*_, …, *x*_35,*i*_) were significantly different from 0, t-test p < 0.01 [corrected]) and the squared error between the averaged fragments of the learning curve in the PD patients and the fragments of the perturbation sequence took its minimal value when the fragments were chosen to be (*p*_9_, …, *p*_32_).

After determining 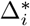 as 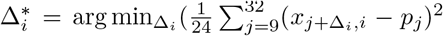, we calculated the amplitude *A_i_* as 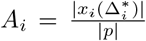 and the phase *ϕ_i_* as 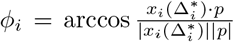, where 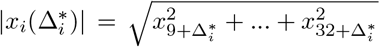 indicates the norm of 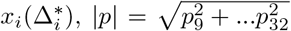 indicates the norm of *p*, and 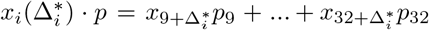 indicates the inner product of 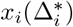 and *p*. Δ_*i*_ indicates the temporal delay with which each subject minimizes the task error according to the imposed perturbation. *A_i_*, the amplitude of learning effects, indicates the response strength to the applied perturbation. *ϕ_i_* indicates the similarity between the learning effects and the applied perturbation sequence. To quantify the similarity between the learning effect and the perturbation while considering the amplitude, the phase, and the temporal delay together, we calculated the root-mean-squared error, 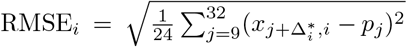. To quantify the trajectories, we calculated the trajectory error as the sum of the lateral deviations within each trial: 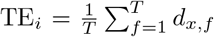, where *d_x,f_* indicates the x-position in the display coordinate system at the *f*th time frame and T indicate the total number of time frames within the trial. We calculated these five variables (i.e., 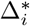, *A_i_*, *ϕi*, RMSE*_i_*, and TE_*i*_) throughout the current study.

### 2.9 Statistical analysis

We utilized one-way ANOVA with a group factor (patients with PD, elderly individuals, and young individuals), followed by Tukey’s post hoc tests to compare each group if there was no specified notification about statistical test. We used MATLAB 2016b (Mathworks, Nantick MA) for all statistical analyses.

## 3 Results

Eighteen PD patients, eighteen age- and MMSE-matched elderly individuals, and eighteen university students (referred to as young individuals in the following) adapted to visuomotor rotation through the PoMLab setting using a tablet device (the age distributions and clinical scores are summarized in Table 1). Figs. 1A and 1B indicate the cursor trajectories and velocities along the y-axis, respectively, in the learning trials averaged across all the subjects in each group. In those figures, the solid green lines indicate participants in the PD patient group, the solid blue lines indicate those in the elderly group, and the solid black lines indicate those in the young group.

We defined the learning effects as the movement angle at the time when the velocity along the y-axis took its peak value in each trial. Because we focused on the adaptation to the visuomotor rotation for which the subjects needed to compensate for the perturbation in the movement angles, the angles were typical values for the discussion of motor adaptation. The movement angles are thus referred to as learning effects hereafter. The angles of visuomotor rotation *p_t_* at the *t*th trial or visuomotor rotation itself are referred to as perturbations hereafter. Although the adaptation to the perturbation can consist of an explicit component (i.e., cognitive ability) and an implicit component (i.e., adaptive component in motor domain) [8,9], we instructed the participants to aim at the target straightforwardly, which enabled us to exclude the explicit components [8,9]. In addition, no participant was aware of the existence of the perturbation, suggesting that the following results mainly consisted of implicit components rather than explicit components.

The learning effects showed between-trial variation depending on the between-trial varying perturbation (Fig. 1C). The shaded area in Fig. 1C denoted the trial numbers (trials 12-35) when the learning effects of PD patients were significantly different from 0 (t-test p<0.01 [corrected]). We compared the learning effects averaged across the trials denoted by the shaded area in each subject (Fig. 1D). There was a significant group effect (F(2,51) = 6.75, p=0.0025), indicating the difference among the PD patients, elderly individuals, and young individuals. There was a significant difference in the learning effects between the PD patients [8.25±0.51, mean±s.e.m., standard error of the mean] and the elderly individuals [6.65±0.31] (p=0.0184) and between the PD patients and the young individuals [6.28±0.36] (p=0.0032). However, there was not a significant difference in the learning effects between the elderly individuals and the young individuals (p=0.8050). In summary, the PD patients had learning effects approximately 20% larger than the those of the elderly individuals and those of the young individuals in these measures.

In contrast to the learning effects, there was no group effect (F(2,51)=1.35, p=0.27) and no significant difference among the three groups (p>0.28) in the RMSE (Fig. 1E), the error between the learning effects and the perturbation (a detailed definition of this metric is in the Methods section). These results indicate that the PD patients showed more substantial learning effects than the elderly individuals and the young individuals but a comparable ability to minimize the RMSE.

To further study the learning effects, we decomposed the learning effects into three components: the amplitude *A*, to quantify the magnitude of the response to the perturbation; the phase *ϕ*, to quantify the similarity between the learning curves and the perturbation; and the temporal delay Δ, to quantify the temporal sensitivity of the response to the perturbation.

For the amplitude measurement (Fig. 2A), there was a significant group effect (F(2,51) = 6.76, p=0.0025), a significant difference between the PD patients [0.910±0.050] and the elderly individuals [0.752±0.025] (p=0.0162), a significant difference between the PD patients and the young individuals [0.721±0.037] (p=0.0034), and no significant difference between the elderly individuals and the young individuals (p=0.842). Fig. 2D shows the learning curves of the representative PD patients who showed large (solid magenta lines) and small (solid cyan lines) amplitude values. The PD patients showed an approximately 20% larger response in the amplitude than the elderly individuals and the young individuals.

**Fig. 2.**
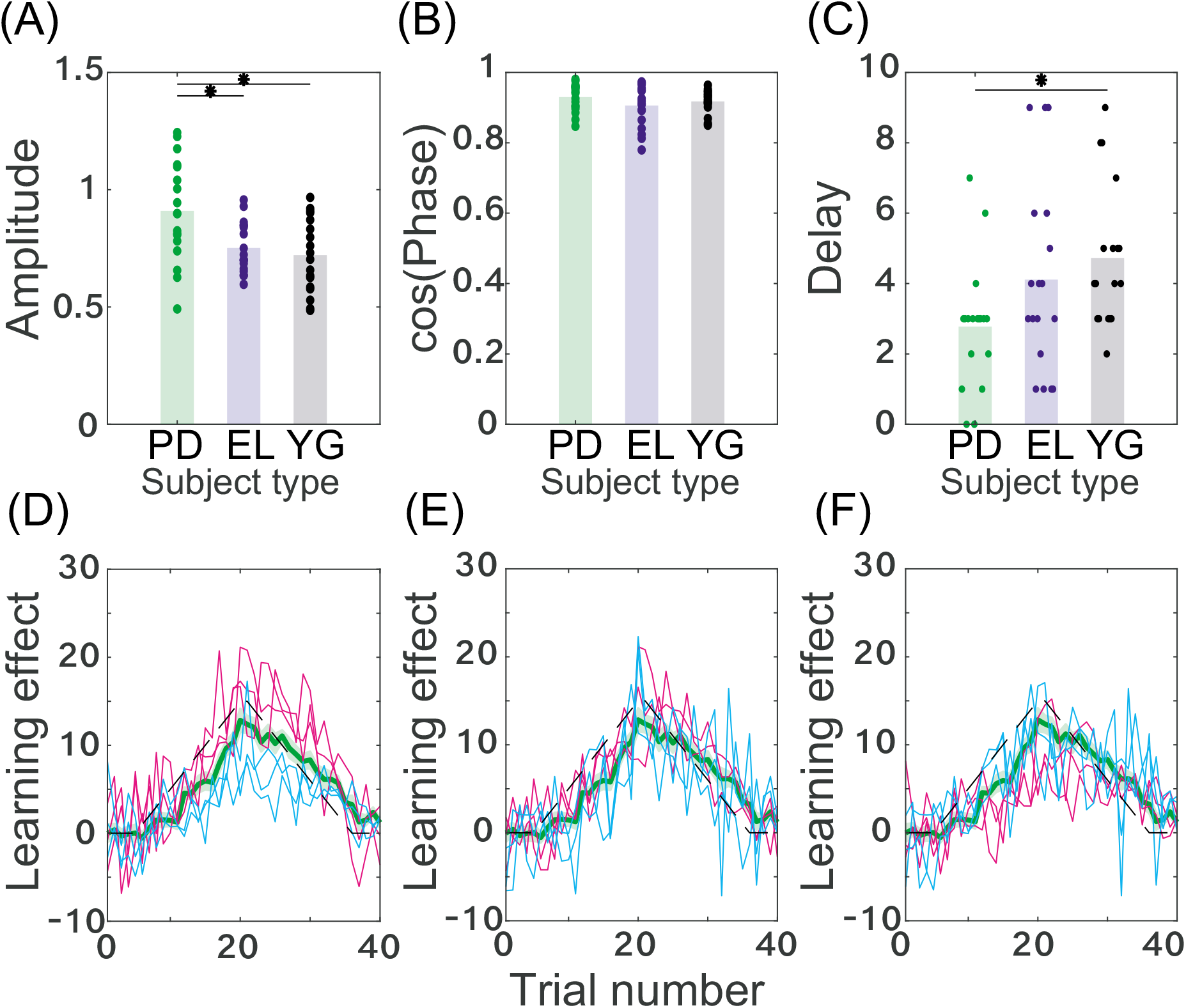
Learning effects decomposed into three factors. (A): Amplitude in each subject and group. A larger value indicates a larger learning effect. Each dot indicates the amplitude for each subject, and each bar shows the mean amplitude in each group. * and ** indicate significant differences with p < 0.05 and p < 0.01, respectively (Tukey’s post hoc test following one-way ANOVA). (B): Cosine function of the phase for each subject and group. A larger value indicate a smaller phase value, which indicates a similar learning curve to the applied perturbation pattern. (C): Delay in each subject and group. A larger value indicate a longer delayed response to the applied perturbation. (D): Typical learning curves in the PD subjects whose amplitude was the largest, the second largest, the third largest (magenta solid lines), the third smallest, the second smallest, and the smallest (cyan solid lines). (E): Typical learning curves of the PD patients regarding the cosine function of the phase. (F): Typical learning curves of the PD patients regarding delays.

For the phase measurement (Fig. 2B), there was no significant group effect (F(2,51) = 1.16, p=0.32) and no significant difference between each group (p>0.3104 among the PD patients [0.930±0.009], the elderly individuals [0.906±0.014], and the young individuals [0.918±0.008]. Fig. 2E shows the learning curves of the representative PD patients who showed large (solid magenta lines) and small (solid cyan lines) phase values. In contrast to the amplitude, the PD patients showed a comparable phase with the elderly individuals and the young individuals.

For the delay (Fig. 2C), there was a significant group effect (F(2,51) = 3.62, p=0.034), no significant difference between the PD patients [2.78±0.42] and the elderly individuals [4.11±0.65], (p=0.18), a significant difference between the PD patients and the young individuals [4.72±0.48] (p=0.030), and no significant difference between the elderly individuals and the young individuals (p=0.688). Fig. 2F shows the learning curves of the representative PD patients who showed large (solid magenta lines) and small (solid cyan lines) of delay values. The PD patients showed smaller response delays compared to the young individuals.

Taken together, the learning effects of the PD patients were larger than the those of the elderly individuals and the young individuals (Figs. 1C and 1D, respectively) because the PD patients showed larger amplitudes compared to the other two groups (Fig. 2A). In addition, a slightly faster response delay in the PD patients contributed to the large learning effects (Fig. 2C).

We further considered other factors that may affect the larger substantial learning effects in PD patients. A candidate in influencing the learning effects was the movement time. Following previous studies that reported slower movement times in PD patients than in elderly individuals [25], those patients also showed slower movement times in our experimental setting than the young individuals (Fig. 3A, significant group effect, F(2,51) = 21.96, p=1.32 × 10^−7^, no significant difference between the PD patients [108.77±6.84 ms] and the elderly individuals [95.77±8.13 ms], p=0.3321, a significant difference between the PD patients and the young individuals [51.41±3.27 ms], p=1.93 × 10^−7^, and a significant difference between the elderly individuals and the young individuals, p=3.11 × 10^−5^). To investigate the possible effects of the movement time on the amplitude, we normalized the movement time within each group so that the mean and the standard deviation of the movement time in each group equaled 0 and 1, respectively. After normalization, we calculated the correlation coefficients between the amplitude and the grouped and normalized movement times. If the movement time affected the learning effects, we could expect some correlation between these metrics. In contrast to this assumption, there was no significant correlation between the magnitude and the normalized movement times (Fig. 3B, r=0.15, p=0.27). Additionally, there was no correlation between the phase and the normalized movement time (r=0.10, p=0.46), between the lag and the normalized movement time (r=−0.15, p=0.28). These results indicate that the movement time was not a significant factor affecting the learning effects.

**Fig. 3.**
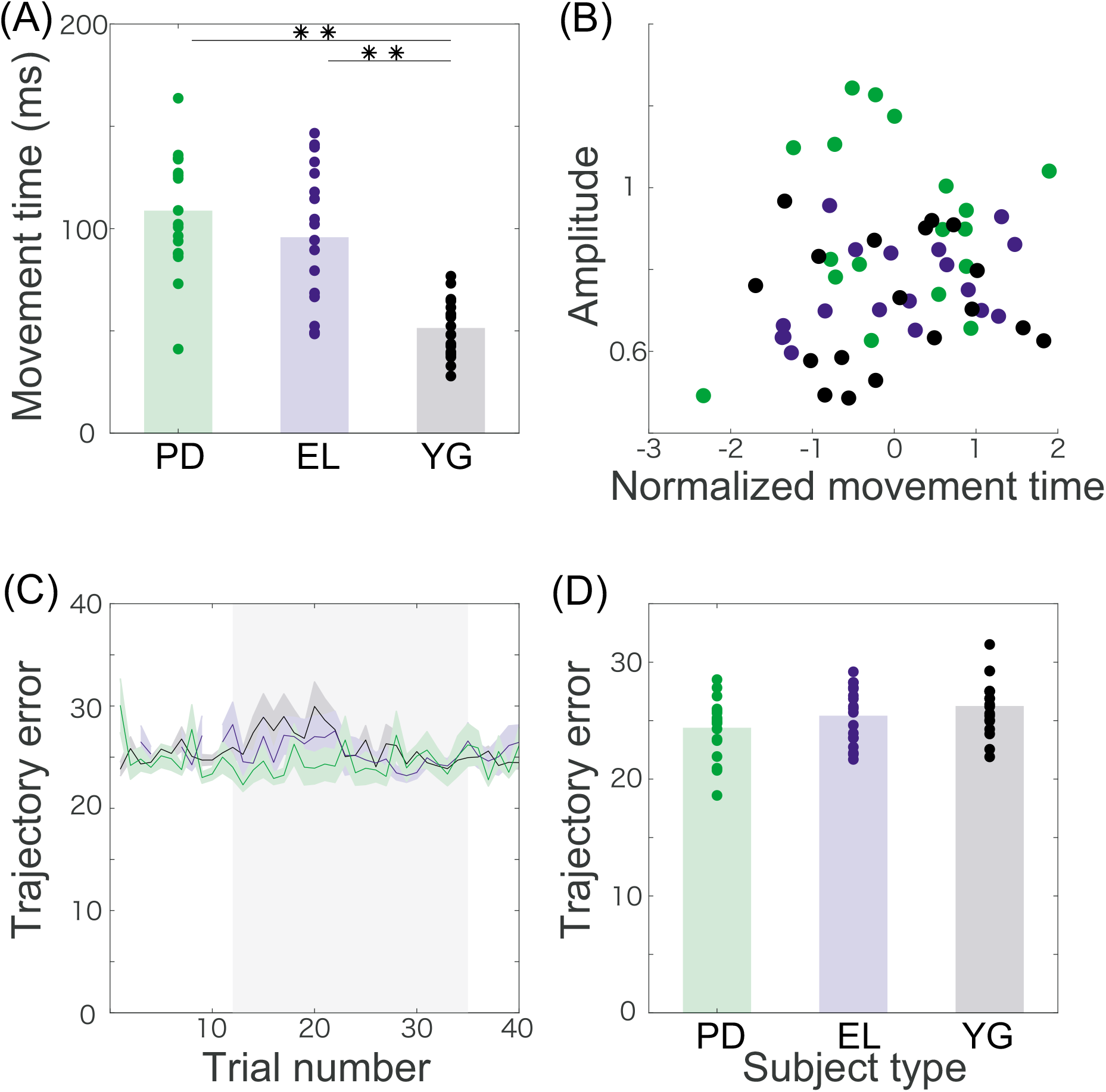
Kinematic factors possibly relating to the learning effects. (A): Movement time. ** indicates a significant difference with p < 0.01 (Tukey’s post hoc test following one way ANOVA). (B): Relation between the normalized movement time and the amplitude of the learning curve. The normalized movement time indicates the modified movement time whose mean and standard deviation are zero and one, respectively, in each group. There was no correlation between the two variables (r=0.1517, p=0.2736). (C): Trajectory error. The horizontal axis indicates the trial number, and the vertical axis indicates the trajectory error. The trajectory error was calculated as the squared lateral deviation of the cursor trajectory. The gray shaded area shows the trial numbers where the learning effects were significantly different from zero (t-test p<0.01 [corrected]). (D): Trajectory error averaged across the trials in the gray shaded area in panel (C). There was no difference among the groups (p>0.0697, Tukey’s post hoc test following one-way ANOVA).

Following previous studies that have reported that PD patients showed a large amount of feedback response [26] and that the feedback response can be a source of motor adaptation [27], we investigated the feedback response as another possible factor affecting the learning effects. Fig. 3C shows the relation, indicated by shaded areas, between the TE during the trials, which is a possible factor reflecting the feedback response, and the learning effects. There was no significant group effect (F(2,51) = 2.58, p=0.086) and no significant difference among the three groups (Figs. 3C and 3D, p=0.43 between the PD patients [24.39±0.60 mm] and the elderly individuals [25.42±0.57 mm], p=0.0697 between the PD patients and the young individuals [26.25±0.57 mm], and p=0.57 between the elderly individuals and the young individuals). These results indicate that the feedback response was not a significant factor affecting the learning effects.

We further investigated the relationship between the clinical scores and the learning effects (Fig. 4). There was no significant correlation between all the recorded attributes and the clinical scores (i.e., age, duration, H&Y, MMSE, and UPDRS) and the properties of the learning effects (i.e., the amplitude and the temporal delay) (Fig. 4, p>0.102 for Pearson’s correlation coefficient, and p>0.208 for Spearman’s rank correlation coefficient). These results indicate that the recorded attributes and conventional clinical scores were not enough to explain the large amplitude and the small response delays in PD patients.

**Fig. 4.**
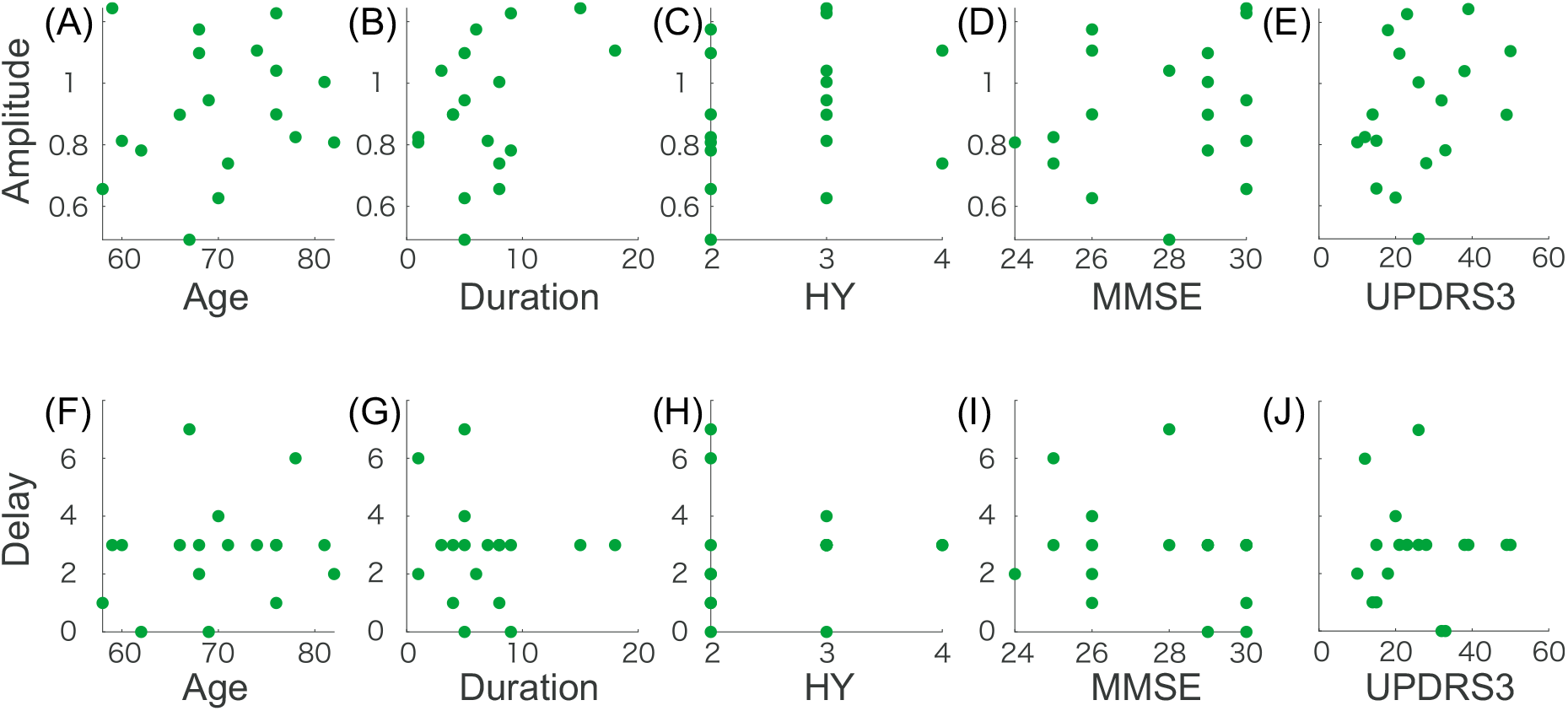
Correlation between the learning effects (amplitude and delay) and the clinical scores of the PD patientso (A-E): The relation between the amplitude and the scoreso (F-J): The relationship between the delay and the scores. There was no significant correlation between the learning effects and the clinical scores (p>0.1024 for Pearson’s correlation coefficient, and p>0.2075 for Spearman’s rank correlation coefficient).

## Discussion

We investigated the motor adaptation ability inherent in PD patients while decreasing the influence of task switching and the burden to participate in the experiments. To reduce the influence of task switching, we utilized a gradual perturbation that was not noticed by most of the participants. To decrease the burden to participate in the experiments, we used a smart-device-based experiment that enabled us to conduct the experiments anytime and anywhere. The current study revealed that the PD patients possessed a motor adaptation ability that was 20% larger, but not better, at minimizing the task errors than the elderly individuals and the young individuals (Figs. 1C-1E and Fig. 2). The larger adaptation ability was not related to the slow movement speed of the PD patients (Figs. 3A & 3B), the feedback response to the perturbation (Figs. 3C & 3D), or the conventional clinical scores (Fig. 4). The larger adaptation ability originated mostly in the larger amplitude (Fig. 2A) and slightly in the faster response delay (Fig. 2C).

A possible factor for the larger motor adaptation ability in the PD patients is compensatory or paretic cerebellar function [28]. The cerebellum plays essential roles in motor adaptation [1,4]. Cerebellar ataxia patients have shown deficits in updating the internal model in motor adaptation experiments [1,4,5]. Although it is widely known that PD patients have paretic symptoms related to dopamine or the basal ganglia, a recent finding supported the possibility that they have a compensatory or paretic cerebellar functions [28]. The modulation of cerebellar function in PD patients can be supported by the connectivity between the cerebellum and basal ganglia [29]. Thus, our findings indicate an aspect of the compensatory or paretic cerebellar function in PD patients, especially in motor adaptation ability.

Another possible factor for the larger observed motor adaptation ability in the PD patients is the reward associated with the motor adaptation task [30,31]. Because the dopamine neurons can encode reward information, such as the temporal difference error [32,33], the lack of dopamine neurons in the PD patients can affect motor adaptation through a deficit in encoding the reward information. In several motor adaptation studies, rewards are associated with accomplishing the performed movements [30,34]. In our experiments, the controlled cursor hit the target before, during, and after adaptation with a high probability (Fig. 1A). Furthermore, in our previous study, we showed that there was no difference between the motor adaptation with and without the vibration in hitting the target in healthy young adults, which was regarded as with and without the reward associated the success of the movement [11]. Thus, we can suggest that our results may originate from compensatory or paretic cerebellar function rather than a deficit in encoding the reward information.

Another possible factor inherent in our findings is the prospective error [18], which is the predicted error in the upcoming movements. In the framework of the prospective error, the predicted error determines a recruitment pattern of neural units responsible for generating motor commands. In this framework, the learning effects when subjects do not predict the prospective error are larger than when they do predict the error. Without the prediction of the prospective error, the same pattern of neural units is recruited across all the trials, and the learning effects are embedded in the pattern in a concentrated manner. With the prediction of the prospective error, a different pattern of neural units is recruited according to the updated prospective error in each trial, and the learning effects are embedded in several patterns in a distributed manner. The learning effects embedded in the concentrated population, rather than in the distributed population, show large learning effects. A possibility inherent in the current finding is that the PD patients possess an impaired ability to predict the prospective error. In some tasks, the PD patients show a deficit prospective ability [35]. This hypothesis can also explain the lack of savings and anterograde interference (a slower learning speed in an interfered task in the A-B paradigm) in PD patients [19,20].

Although the state-space model [36–39] is a popular method to quantify the learning effects in motor adaptation, it is not appropriate in the current study. In this framework, the task error should be minimized. In our current setting, the learning effects of PD patients took smaller values than the perturbation sequence until the 24th trial and larger values from the 25th trial. In this case, the framework of the state-space model predicted that the learning effects increased in each trial until the 24th trial; however, in our experimental setting, the leaning effects decreased from the 20th trial. Thus, we did not apply the state-space model in our current setting. Our findings suggest the need to improve the state-space model to explain adaptation to gradually applied and vanishing perturbations.

Promising future work using PoMLab would be to investigate motor adaptation ability inherent in several types of patients, such as stroke patients, cerebellar ataxia patients, and Huntington disease patients, or the ability associated with autism spectral disorder, schizophrenia, etc. A cross-syndrome comparison can provide essential knowledge about the neural mechanisms of updating the internal model. Although several studies have investigated the ability of patients [4,5,26,40], the experimental setting is different in each study. PoMLab is a cross-platform application and is available for free on our GitHub page (https://github.com/masahiroshinya/PoMLab), which can help researchers, physical therapists, medical doctors, or anyone conduct motor adaptation experiments. Additionally, PoMLab supports conducting motor adaptation experiments anytime and anywhere while decreasing the burden to participate in an experiment.

## Acknowledgments

We thank the staff at Kakeyu Rehabilitation Hospital for their kind support for this research. In particular, we thank Youichi Maruyama and the physical therapists at the hospital. We also thank Shin-ichi Muramatsu for his help and Shintaro Uehara for his helpful comments.

We acknowledge support from the Nakajima Foundation, the Kayamori Foundation of Informational Science, a Novartis Pharma Research Grant, and Promoting Science (a Grant-in-Aid for Young Scientists (18K17894), and Grant-in-Aid for Challenging Exploratory Research (16K12988)).

## Additional information

Competing interests: The authors declare no competing financial or nonfinancial interests.

## Data availability

The datasets analyzed in the current study are available from the corresponding author upon reasonable request.

## Author contributions

K.T. and M.S. designed the experiments and T.S., T.S., and H.O. performed the experiments. K.T. performed the analyses and wrote the manuscript. T.K. oversaw the manuscript.

